# Atomistic Insights Into The Mechanism of Dual Affinity Switching In Plant Nitrate Transporter NRT1.1

**DOI:** 10.1101/2022.10.17.512638

**Authors:** Balaji Selvam, Jiangyan Feng, Diwakar Shukla

## Abstract

Improving nitrogen use efficiency is critical to enhancing agricultural productivity and to mitigate environmental pollution. To overcome the fluctuations in soil nitrate concentration, plants have evolved an elaborate nitrate transporting mechanism that switches between high and low affinity. In plants, NRT1.1, a root-associated nitrate transporter, switches its affinity upon phosphorylation at Thr101. However, the molecular basis of this unique functional behavior known as dual-affinity switching remains elusive. Crystal structures of the NRT1.1 nitrate transporter have provided evidence for the two competing hypotheses to explain the origin of dual-affinity switching. It is not known how the interplay between transporter phosphorylation and dimerization regulates the affinity switching. To reconcile the different hypotheses, we have performed extensive simulations of nitrate transporter in conjunction with Markov state models to elucidate the molecular origin for a dual-affinity switching mechanism. Simulations of monomeric transporter reveal that phosphorylation stabilizes the outward-facing state and accelerates dynamical transitions for facilitating transport. On the other hand, phosphorylation of the transporter dimer decouples dynamic motions of dimer into independent monomers and thus facilitates substrate transport. Therefore, the phosphorylation-induced enhancement of substrate transport and dimer decoupling not only reconcile the competing experimental results but also provide an atomistic view of how nitrate transport is regulated in plants.

## INTRODUCTION

Nitrogen (N) is an essential nutrient for plant growth, development and reproduction.^1^ Plants source nitrogen mostly in the form of synthetic fertilizers where ∼110 million tons of nitrogenous fertilizers are used annually to maximize crop yield.^2,3^ However, plants only use 25-50% of the applied nitrogen, so any excess soil nitrogen results in biodiversity loss, environmental pollution, and contributes to climate change. The global population is projected to increase to ∼10 billion by 2050 and maintaining food supply to feed the world population requires modern technologies to enhance productivity in agricultural lands.^4^ Effective nitrogen management could potentially increase the crop yield and minimize the nitrogen loss to the environment. Improving the nitrogen use efficiency (NUE) will allow sustainable and eco-friendly food production.^2^ It is estimated that a 1% increase in NUE could save ≥ 1.1 billion US dollars each year.^2^

Being immobile, plants acquire various nutrients from soil through root-associated transporter proteins in the plasma membrane. For most terrestrial plants, nitrate is the preferred nitrogen source and is readily absorbed by plants at the root surface via nitrate transporters. Biochemical studies have shown that these transporter proteins can adapt to fluctuations in the soil nitrate concentration. To ensure sufficient nitrate supply even under highly fluctuating environment, plants have evolved low-affinity (LATS) and high-affinity (HATS) nitrate transporter systems.^5^ Intriguingly, the nitrate transporter NRT1.1 (also known as CHL1 or NPF6.3) in *Arabidopsis thaliana* is a dual-affinity transporter, which switches between low and high affinity modes in response to changes in the soil nitrate concentration through phosphorylation of an intracellular threonine residue Thr101, localized at the N-terminal end of transmembrane helix 3 (TM3).^6–9^ At low nitrate concentration, Calcineurin-B Like (CBL)-Interacting serine/threonine Protein Kinase (CIPK23) phosphorylates NRT1.1 at Thr101, which increases the affinity for the nitrate ion (K_M_ ≈ 50 µM) and results in a high affinity transporter. When nitrate is abundant, NRT1.1 is not phosphorylated at Thr101 and acts as a low affinity transporter (K_M_ ≈ 4 mM). This inherent regulatory mechanism of NRT1.1 helps plants to adapt and respond to varying nitrate levels in the soil. However, the mechanistic basis of nitrate uptake and structural mechanism of dual-affinity switching remains elusive.

NRT1.1 belongs to the proton-dependent oligopeptide transporter family (POT) of secondary active transporters.^10^ Despite the low sequence similarity, proteins in POT family share a conserved structural topology with major facilitator superfamily (MFS) of transporters and are considered as a distant member of the MFS. NRT1.1 adopts the canonical MFS fold, with TM1-6 and TM7-12 forming the N and C-terminal bundle, respectively. MFS family membrane transporters undergo a cycle of conformation change from an outward-facing (OF) state to occluded (OC; both extracellular and intracellular gates are closed) and finally to the inward-facing (IF) state.^11^ Recently, two independent studies have reported the X-ray crystal structures of NRT1.1 in an almost-identical IF conformation and have provided the first glimpse of complex molecular interplay between transporter activity and post-translation modifications.^12,13^ The nitrate ion was characterized at the center of transporter pore tunnel and stabilized by His356 at the binding site.^12,13^ NRT1.1 was crystallized as a dimer with both the monomers in the IF conformation with Thr101 located close to the dimer interface. Cell-based fluorescence and *in vitro* biochemical studies have shown that transient dimer of NRT1.1 could be formed in both cell and detergent solutions. Based on these observations, Sun *et al*. propose that phosphorylation of Thr101 affects the dimerization of NRT1.1. However, it is not clear how dimerization affects the nitrate K_M_ value. Parker and Newstead have questioned the regulatory role of NRT1.1 dimerization and suggested that it could be a crystallographic artifact. The Thr101Asp mutant of NRT1.1 (a change that is used as a phosphomimic) have the same K_M_ value as wild-type NRT1.1, much higher transport rate, and a lower melting temperature. These observations indicate that Thr101Asp mutant has a higher transport rate due to enhanced structural flexibility. To reconcile these disparate views, a dimerization switch model has been proposed -- dimer decoupling, induced by phosphorylation, increases the structural flexibility.^8^ However, direct experimental or computational evidence for the model has not been reported.

From these structural and biochemical studies, the dimerization, Thr101 phosphorylation and the structural plasticity of the transporter are proposed to play a key role in the affinity switching and the substrate translocation mechanism of NRT1.1. However, many important questions remain unanswered: How does dimerization and phosphorylation tune the substrate affinity? Is a dimerization-switch model valid? How does phosphorylation affect the transport properties of NRT1.1 monomer? What is the physiological relevance of NRT1.1 dimerization and what are the cascade effects of phosphorylation on NRT1.1 dimer? And finally, what is the molecular mechanism of the nitrate transport coupled conformational transitions in NRT1.1?

To address these unanswered questions, we have performed extensive molecular dynamics simulations to explore the conformational dynamics and nitrate transport mechanisms of the Unphosphorylated (UnpNRT1.1) and phosphorylated (pNRT1.1) forms of the AtNRT1.1 monomer and dimer. MD simulations have been successfully over the last three decades used to decipher the impact of post-translational modifications on protein dynamics and function.^14^ Using Markov state models (MSMs), we have characterized the complete nitrate transport cycle in both UnpNRT1.1 and pNRT1.1. Our simulations show that phosphorylation increases structural dynamics of the N and C-terminal domains and thus increases the substrate transport. Addition of the negatively charged phosphate group on Thr101 results in partial helical unwinding at the bottom of TM3, which allows the adjacent positively charged residues to form favorable interactions, thereby accelerating the structural transition from the IF to OF state. To further investigate the phosphorylation-induced NRT1.1 dimer dissociation, we employed umbrella sampling simulations for both the unphosphorylated and phosphorylated dimers. These simulations revealed a high free energy barrier during the conformational transition from IF to OF in UnpNRT1.1 dimer as compared to the pNRT1.1. In other words, dimerization restricts the dynamic motions of each individual monomer thereby reducing transport rate, whereas the Thr101 phosphorylation decouples the dimer and increases transport rate. Finally, these simulations also provide the relative population of the different conformational states which explains how phosphorylation-induced dimer decoupling regulates the K_M_ of NRT1.1.

## METHODS

### Molecular dynamics (MD) simulation details for UnpNRT1.1 and pNRT1.1 monomer

All MD simulations were performed using AMBER14.^15^ The default force field AMBER FF14SB force field was used for the simulation.^16^ The nitrate ion and phosphorylated threonine parameters were obtained from AMBER parameter database. The simulation systems were set up using the tleap program in AmberTools14. The starting coordinates of NRT1.1 was obtained from Protein Data Bank (PDB) (PDB ID: 4OH3).^17^ The chain termini were capped with neutral acetyl and methylamide groups. The structure was embedded in a phosphatidylcholine (POPC) bilayer and solvated in orthorhombic boxes with dimension of 87 Å * 87 Å * 109 Å with TIP3P water molecules.^18^ The MD system was neutralized by adding 0.15 M of salt concentration. The titratable residues were assigned to default protonation state. The histidine residue His356 was protonated by assigning net charge of +1.^12,13^ All simulations were performed with protonated histidine. Threonine residue Thr101, the known phosphorylation site in NRT1.1, was phosphorylated and the MD system was referred as phosphorylated NRT1.1 (pNRT1.1) simulation and the same was kept unphosphorylated in Unphosphorylated NRT1.1 (UnpNRT1.1) simulation. The final MD systems of both UnpNRT1.1 and pNRT1.1 contain ∼85,000 atoms. The MD system was minimized using conjugate gradient method for 20,000 steps and slowly heated from 0 to 10 K, and then 10 to 300 K over a period of 1 ns each in canonical (NVT) and isothermal-isobaric (NPT) ensembles. Both the systems were equilibrated in an NPT ensemble for 50 ns and the production runs were carried out in the NPT ensemble at 300 K and 1 atm. Previous work of ours has shown that use of the different thermostat and barostat maintains free energy landscape integrity.^19^ Periodic boundary conditions (PBC) were applied to all simulations, particle mesh Ewald (PME) was used to treat long-range non-bonded interactions.^20^ The SHAKE algorithm was used to constrain hydrogen-containing bonds. A 2 fs time step was used throughout all simulations.^21^ To generate a diverse set of initial structures for conventional MD (cMD) simulations, we performed accelerated MD (aMD) for both UnpNRT1.1 and pNRT1.1. A boost potential of 0.8 and 0.2 were used for all dihedrals and for the entire MD systems, respectively. The total duration of aMD for UnpNRT1.1 and pNRT1.1 is ∼24 and ∼11 µs, respectively. The aMD simulation data were clustered based on the extracellular and intracellular gating residue pairs and 200 clusters for both the MD system was obtained. The snapshots from the cluster center was extracted and used as a starting structure for the cMD. Adaptive sampling approach was used to capture the rare transition events.^22^ Trajectory snapshots were collected every 100 ps. A total of ∼142 µs and ∼63 µs was obtained for both UnpNRT1.1 and pNRT1.1, respectively.

### Markov state model construction and trajectory analysis

The generation of Markov state models (MSMs) is a powerful method to characterize the kinetic and thermodynamic properties of complex biomolecular systems. MSM constructs a transition probability matrix between a set of microstates estimated from a large number of short independent MD trajectories.^23–26^ MSMs have been recently used to investigate the conformational dynamics associated with membrane transporters.^19,24,27–31^ MSMs were constructed using MSMBuilder 3.6.^32^ The simulation data was analyzed using MDTraj ^33^. The MD snapshots were visualized using VMD1.9.2 and pymol 2.^34,35^ The data from cMD simulations were used for MSM analysis and the Markovian lagtime for each MSM was chosen from the implied timescales plot (Fig. S8). The first and most critical step in MSMs construction is featurization, projecting high-dimensional MD data into a low-dimensional feature space. A subset of alpha carbon distances between residues were chosen as input features and further coarse-grained using the time-lagged independent component analysis (tICA) technique.^36^ tICA was developed to obtain the slowest collective motions by linear combinations of input degrees of freedom. The featurized MD data was clustered into conformational microstates with the mini-batch K-means clustering algorithm.^37^ The distance features and the hyperparameters for MSM construction are systematically chosen using genetic algorithm. The top flux pathways for the conformational transition between the intermediate states were obtained using TPT and the kinetic properties were determined using Mean First Passage Time (MFPT). MFPT identifies the mean transition time from one state to another state. The free energy for nitrate at different states was calculated from a one-dimensional potential of mean force (PMF). The dynamic coupling plots were obtained using MD-TASK. The pore radius plots were obtained using HOLE program.^38^ All the plots were generated using MATLAB_R2015b.

### Genetic algorithm

To select the “best” MSM automatically, we combined the genetic algorithm and Osprey variational cross-validation package to optimize the set of Cα atom distances between residue pairs along with two critical hyperparameters (number of tICA components and number of clusters) in MSM construction.^39^ The quality of MSMs is quantified with the generalized matrix Raleigh quotient (GMRQ).^40^ GMRQ is the sum of the eigenvalues of the transition matrix, indicating that the higher the GMRQ, the better the MSM. For each set of distances, 10 MSMs were built by (1) featurizing MD data based on the set of distances, (2) decomposing the featurized data using tICA, (3) and then clustering the data with the mini-batch K-means clustering algorithm. The two hyperparameters (number of tICA components and number of clusters) were varied in the 10 MSMs. Finally, GMRQ score was assigned to each MSM and the best GRMQ score of the 10 MSMs was used as the “fitness” of the set of distances. The GRMQ score varying with iterations of the genetic algorithm is shown in Fig. S9. Eventually, the MSM with the highest GMRQ among all the iterations was chosen for further analysis (detailed in Table S1).

### Umbrella sampling protocol for UnpNRT1.1 and pNRT1.1 dimer

To investigate the role of phosphorylation in NRT1.1 dimerization, we employed umbrella sampling to simulate the conformational dynamics of UnpNRT1.1 (∼4 µs) and pNRT1.1 (∼4 µs) dimer. We selected the distance root-mean-square displacement (dRMSD) with respect to the starting crystal structure (PDB: 4OH3) as collective variables. Two collective variables are defined as the dRMSD calculated at the extracellular and intracellular helix tip region, respectively. Because these two collective variables strictly determine the IF, OC and OF, respectively. The force constant of the harmonic biasing potential was 3000 kcal/mol*nm^2^ for all reaction coordinates. To generate starting structures for umbrella sampling, cMD was employed for both the UnpNRT1.1 (∼9 µs) and pNRT1.1 dimers (∼5 µs). The initial umbrella sampling windows were created every 0.5 Å in each dimension. Since the starting configures are either in the inward-facing or the partial inward-facing occluded state, we applied adaptive umbrella sampling where a set of new windows were selected from the previous round of sampling.^41^ In total, 291 and 340 windows were generated for UnpNRT1.1 and pNRT1.1 dimer, respectively. Each window was simulated for ∼15 ns. The PLUMED2.4.2 package was used to patch GROMACS 5.1.4 and define the collective variables for biasing.^42^ All umbrella sampling simulations were performed in GROMACS 5.1.4, using a time step of 2 fs.^43^ Semi-isotropic Parrinello-Rahman pressure coupling with a reference of 1 bar in both XY and Z directions was used for production simulations. The Nose-Hoover thermostat was used for production with time constant of 0.5 ps and reference temperature of 300 K. Coulombic interactions were calculated using the Particle Mesh Ewald (PME) method, with a grid spacing of 0.16 nm.^20^

## RESULTS AND DISCUSSION

### Thr101 phosphorylation stabilizes the outward-facing state in monomeric transport to enhance substrate affinity

The inward-facing (IF) crystal structure (PDB ID: 4OH3) was used as starting structure for the MD simulations. Thr101 was converted to phosphorylated Thr101 and simulations were performed over a duration of ∼142 µs and ∼63 µs for the UnpNRT1.1 and pNRT1.1 monomers to explore their conformational dynamics and characterize the complete nitrate transport mechanism. Markov state models (MSMs) were constructed using the simulation datasets by clustering along the five slowest timescales process (see Methods for details) and the final model contains 200 conformational states. MSMs allow the representation of the simulation data as conformational states of the protein and the rates between these states. This network representation can be used to estimate the thermodynamic and kinetic properties such as free energy or stability for different transporter conformation and the rates of substrate transport.

An MSM weighted conformational free energy landscape plot was generated by projecting the simulation data along the extracellular and intracellular minimum helix tip distances, as they show significant correlation with minimum tunnel radius that substrates can pass through for each of the gates.^44^ The conformational dynamics reveal that UnpNRT1.1 exhibits more diverse conformational states; in contrast, pNRT1.1 follows the canonical L-shaped landscape to resemble an alternate-access model (Fig.1). The calculation of helix tip distance of N- and C-terminal domain of the IF structure reveals that the extracellular helices are packed closely to each other and the intracellular ends are open with a minimum distance of ∼5 Å and ∼15 Å, respectively. The formation of polar contact between Lys164 (NZ)-Tyr480 (CZ) initiates the structural transition to other intermediate states. As the distance decreases to ∼3-4 Å, the helices 4-5 and 10-11 approach one another to close the intracellular end which leads to the OC state. Residues Arg183 (TMH5) and As485 (TMH10) form ionic contact which in turn stabilizes the OC state. The OC state is relatively stable in both UnpNRT1.1 and pNRT1.1 and the free energy barrier in the order of less than 0.5 kcal/mol was estimated for the conformational transition from IF to OC states. The stronger contacts at the intracellular domain destabilize the hydrophobic contacts at the extracellular side and initiates the structural rearrangement to the OF state. The OF state is easily accessible in pNRT1.1 whereas a high energy barrier is noticed for UnpNRT1.1. The extracellular helical tips move far apart by up to ∼14 Å to reach the OF state. The conformational landscape plot reveals that OC to OF transition in pNRT1.1 has a free energy barrier of ≤1.5 kcal/mol. Our data then supports that phosphorylation enhances the structural plasticity of the transporter, thereby stabilizing the OF state. However, OF is identified as a high-energy state in UnpNRT1.1 with a free energy barrier of ∼ ≥2.5 kcal/mol. In other words, the transition to OF is an energy-demanding process, and these states are the least stable in UnpNRT1.1.

**Fig. 1.**
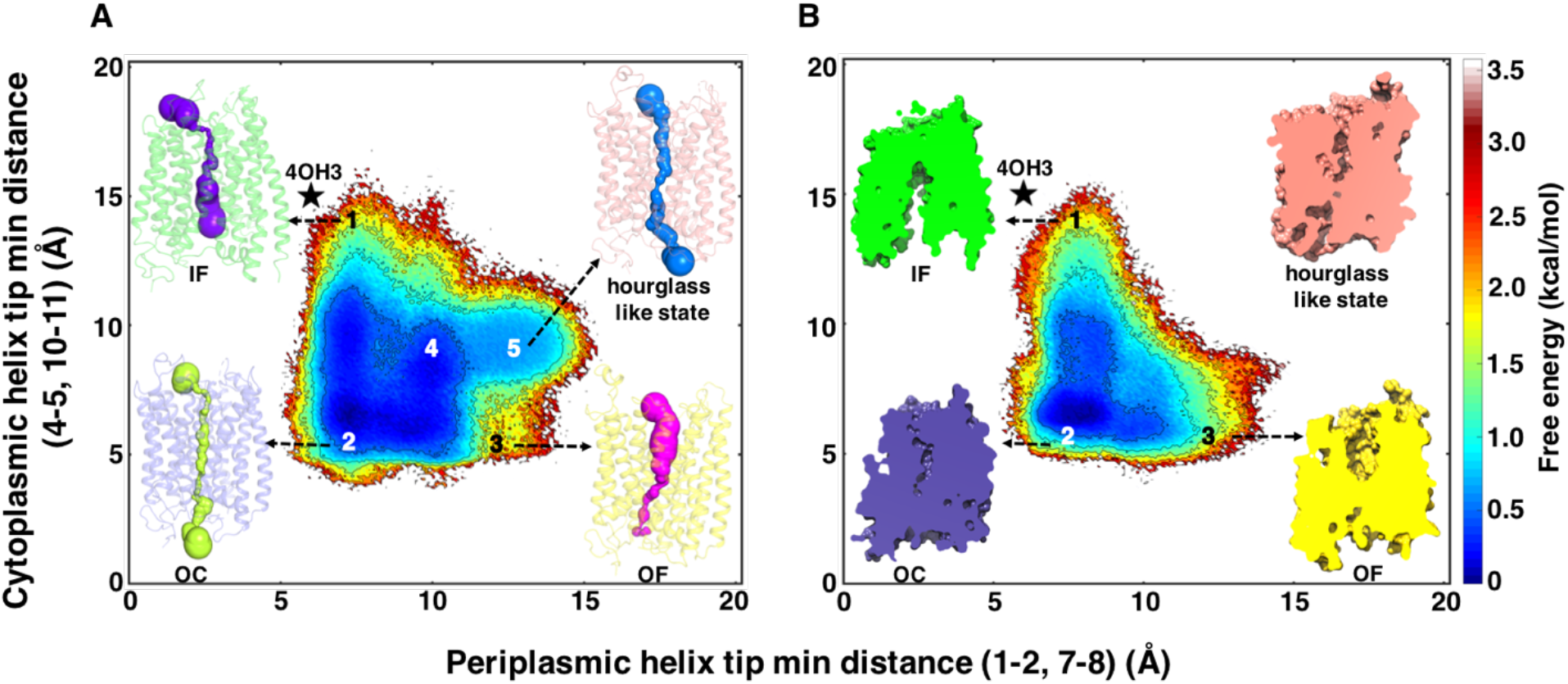
Conformational dynamics of (A) UnpNRT1.1 and (B) pNRT1.1 monomer. The opening and closure of channel gates are measured by the minimum helical tip distance of first four residues at the helices 1-2 and 7-8 at the periplasmic side and the helices 4-5 and 10-11 at the cytoplasmic side. The intermediate states are depicted as (1) IF, (2) OC, (3) OF, (4) partial IF-OF, and (5) hourglass-like state, respectively. The pore radius of the transporter channel for intermediate states was calculated using the HOLE program^38^ and represented as a surface (Fig. 1A). The residues that drive the structural rearrangements and that stabilize the metastable conformational states are shown in ball and sticks (Fig. 1B). TPT shows two dominant pathways for UnpNRT1.1 from IF to OF via OC (1→2→3) and through partial IF-OF (1→4→3). For pNRT1.1, one dominant pathway was observed from IF to OF via OC (1→2→3).

The landscape plot also shows that UnpNRT1.1 samples a diverse conformational space such as partial IF-OF and hourglass-like states. The pore channel radius plot reveals that the transporter is open at both ends and constricted at the center of the pore channel in these states. Membrane transporters adopting hourglass-like topology have been identified in bacterial acetate transporter and its biological implications in substrate translocation has been demonstrated using MD simulations^27,45^. To further investigate the role of these intermediate states, we used transition path theory (TPT) to identify the dominant pathways of conformational transition from IF to OF (Fig. S1).^46^ TPT identifies the highest flux paths and provides the probability of the most preferred pathway for the structural transition to various intermediate states. The kinetic data estimated from MSMs reveals that pNRT1.1 monomer transports nitrate faster than UnpNRT1.1 monomer (Fig. S2). The average time required for one complete cycle of nitrate translocation from IF to OF in pNRT1.1 and UnpNRT1.1 monomers was predicted as 5 ± 2 µs and 18 ± 3 µs, respectively (Fig. S2). Our study suggests that phosphorylation changes the conformational dynamics and leads to fast conformational transition from IF to OF via OC by reducing the stability of other intermediate states while stabilizing the OF state.

### Thr101 phosphorylation enhances structural flexibility and dynamic coupling between N and C domains of the monomeric transporter

To gain insights into the effect of Thr101 phosphorylation (pThr101) on structural and dynamic changes, we examined the difference of charge distribution on the surface of different states between UnpNRT1.1 (Fig. 2A) and pNRT1.1(Fig. 2B). The electrostatic-potential maps show that pNRT1.1 undergoes dramatic changes in the charge distribution compared to UnpNRT1.1. In the IF crystal structure, Thr101 is buried inside the helical bundle surrounded by hydrophobic residues (Ile91, Leu96, Leu100, Ile104 and Phe105) at the intracellular end of helix 3. A similar trend was observed in our simulation and the charge density maps show that Thr101 is buried in a hydrophobic pocket in OC and OF state (Fig. S4). The addition of a phosphate group in Thr101 increases the positively charged patch at the surface of the intracellular side of helix 3. The accumulation of negative charges causes helical unwinding which results in twisting of the loop linking residues at TMH2 and TMH3. This exposes the backbone atoms of Gly97, Arg98, Tyr99 and Leu100 to form polar contact with pThr101 to neutralize the excess negative charges. In UnpNRT1.1, Arg98 interacts with Asp173 from TMH4 in IF, OC and OF states, whereas Arg98 interacts strongly with pThr101 and Asp93 in OC and OF states. The charge-charge induced electrostatic changes lead to structural rearrangement in the N-terminal domain and partially close the intracellular half of NRT1.1, hence facilitating the conformational transition from IF to OC (Fig. S4). The computed solvent-accessible surface area (SASA) of residues around the vicinity of phosphorylated residue in UnpNRT1.1 and pNRT1.1 indicate that Thr101 is buried in a hydrophobic pocket and gets exposed to solvent upon phosphorylation (Fig. S3). To further understand the role of phosphorylation induced fast conformational transitions, we performed cross-correlation analysis of inter-domain motions in both UnpNRT1.1 and pNRT1.1 simulation data (Fig. S5). The motions of N-terminal and C-terminal domains are positively correlated and dynamically coupled in pNRT1.1. The high degree of correlated movements was observed between helix 2-3 and C-terminal domain, in contrast, the respective regions are less coupled and negatively correlated in UnpNRT1.1. The correlation plots also reveal that the dynamics of helices 4-5 exhibit strong correlation with helices 7-8, 9 and 11 of pNRT1.1 and is involved in allosteric communication between distal residues. The loosely coupled nature of these specific domains prevents the efficient functioning of UnpNRT1.1 and results in a low affinity transporter. In summary, phosphorylation triggers the helical unwinding at the intracellular side of helix 3, which in turn enhances the dynamic coupling between N and C-terminal domain to drive active transport.

**Fig. 2.**
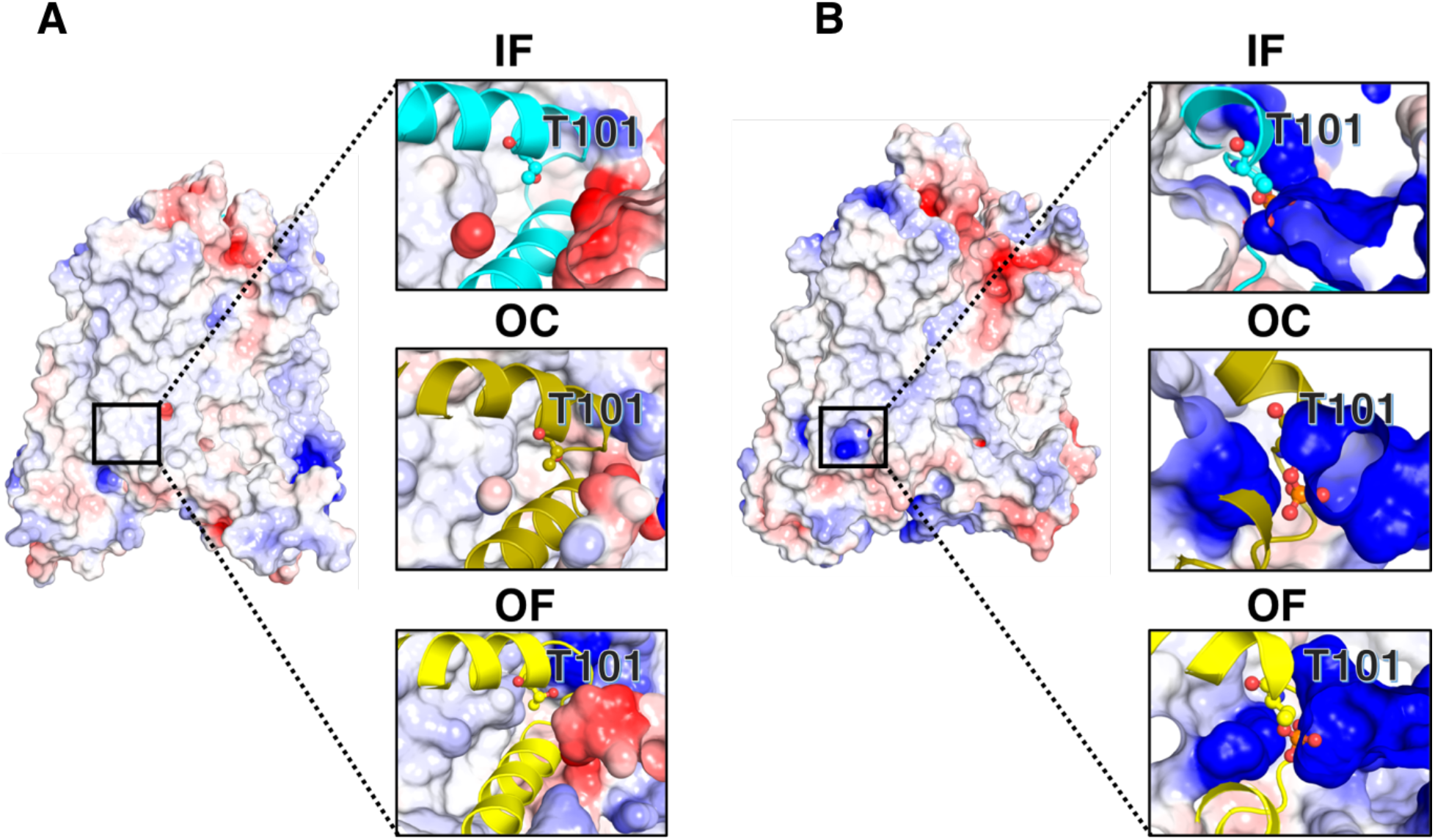
Role of phosphorylation. The charge distribution surface around the **(A)** unphosphorylated and **(B)** phosphorylated Thr101 region was shown for IF, OC and OF states with an electrostatic scale from -5 to +5 eV corresponding to red and blue colors.

### Phosphorylation decouples the dynamics of the NRT1.1 dimer

Besides altering the transport properties of the NRT1.1 monomer, phosphorylation also plays a role in NRT1.1 dimerization. To examine the biological relevance of dimerization and its effect on structural flexibility of NRT1.1, we employed umbrella sampling for both UnpNRT1.1 and pNRT1.1 dimer (see Materials and Methods for details). Umbrella sampling is an enhanced sampling method that uses a bias potential to enable conformational characterization of large systems such as NRT1.1 dimer. Using distance root-mean-square displacement (dRMSD) with respect to the crystal structure as a collective variable, we obtained ∼4 µs of simulation data for both the dimer systems. The simulation data was projected along the extracellular and intracellular helix tip minimum distance root mean squared deviation (dRMSD) to characterize the conformational transition to intermediate states. The potential of mean force (PMF) plot reveals that a high free energy barrier on the order of ∼20-23 kcal/mol is encountered in UnpNRT1.1 dimer for the structural transition to OF via the OC state (Fig. 3). However, the OF state is relatively more accessible in pNRT1.1 dimer and the conformational dynamics plot reveals that free energy barrier is estimated as ∼8-12 kcal/mol. The resolved crystal structure shows that the monomers are positioned adjacent to each other with an extensive surface area of ∼2160 Å^2^ in UnpNRT1.1 dimer and significantly affect the overall dynamics of the complex. The landscape plot also reveals that transition to OC state is favorable in pNRT1.1 (∼3 kcal/mol) while achieving the OC state is still an energetically demanding process in UnpNRT1.1 (∼10 kcal/mol). To further investigate the specific interactions between monomers in UnpNRT1.1 and pNRT1.1 dimers, we performed unbiased simulation over a period of ∼9 µs and ∼5 µs, respectively. Cross-correlation analysis on this data shows that direct coupling at the dimeric interface across the monomers were identified between the helices 3-6 in UnpNRT1.1 dimer, whereas monomers behave independently in pNRT1.1 dimer (Fig. S6). Our simulation results are in agreement with previous fluorescence resonance energy transfer (FRET) results that the dynamical motions are coupled in UnpNRT1.1 dimer and phosphorylation uncouples the motion between the two monomers.^12^ Thus, phosphorylation destabilizes the dimer and enhances the transport rate by allowing higher structural flexibility in each monomer (Fig. S6C).

**Fig. 3.**
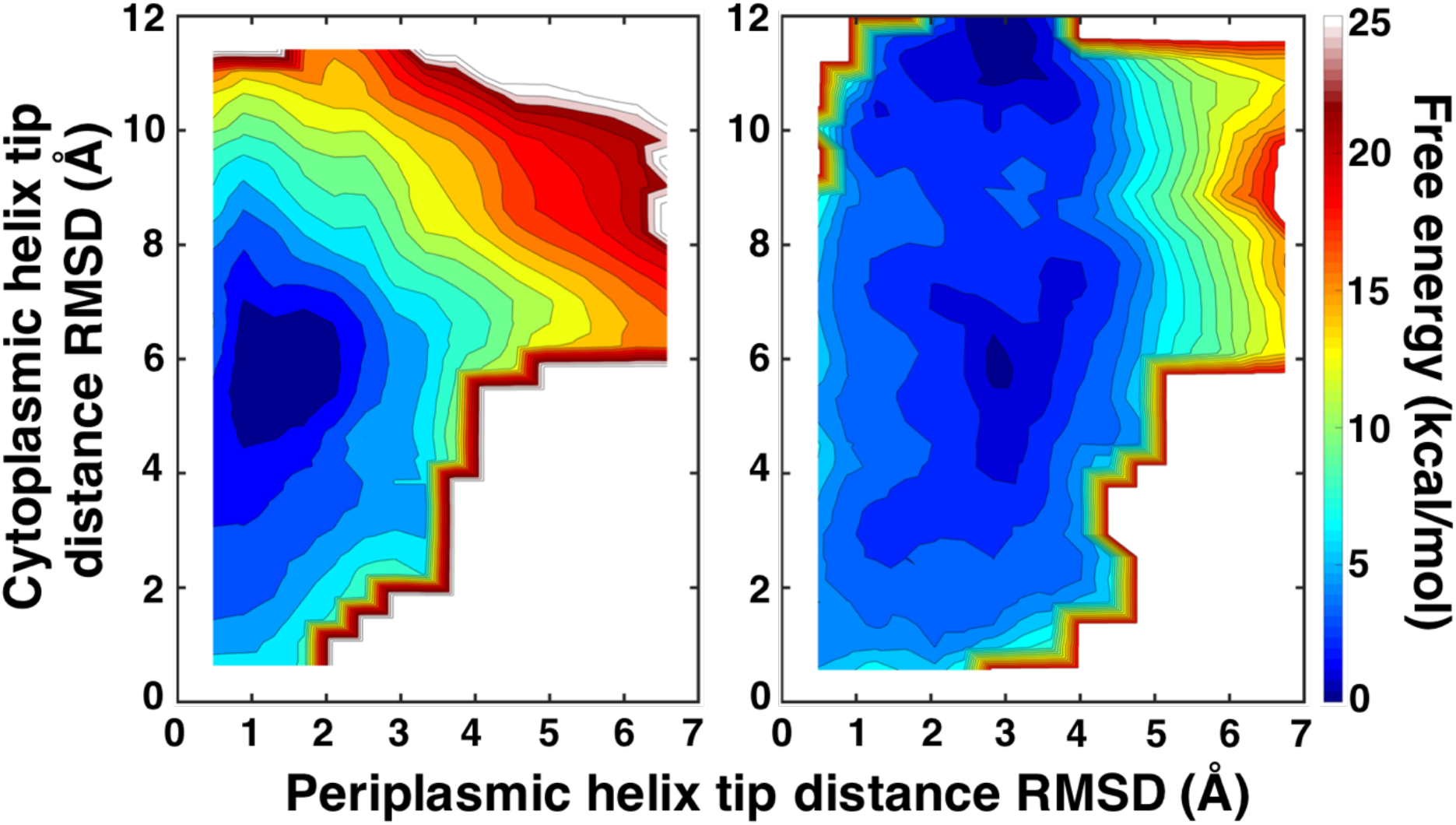
Conformational dynamics of UnpNRT1.1 and pNRT1.1 dimers. Free energy landscapes of **(A)** UnpNRT1.1 and **(B)** pNRT1.1 dimers are projected along the distance root mean square deviation (dRMSD) of the periplasmic and cytoplasmic helix tip in reference to the crystal structure (PDB: 4OH3).

### Simulations characterize the nitrate recognition, binding, and translocation cycle

To understand the molecular mechanism of nitrate recognition, binding and translocation, we extracted the ensemble of structures from the individual states of highest flux path as determined by TPT (Fig. 4). The nitrate ion is recognized by positively charged residues Arg129 and Arg217 at the extracellular surface (Fig. 4A). In the OF state, the distances between the extracellular and intracellular gating residues are ∼13.5 Å and ∼4 Å, respectively. The ion enters the translocation pore and forms polar contact with Thr360 (Fig. 4B). Nitrate escapes from the intermediate interaction and diffuses further inside to bind to the protonated His356 on TMH7, in accordance with the experimental finding that the mutation of His356 to alanine abolishes nitrate uptake as well as transport in both low and high-affinity mode.^12,13^ The rotameric conformation of His356 is further favored by Glu476 and Tyr388 as these residues interact together (His356-Glu476-Tyr388) and form a polar contact triad (Fig. 4C). Arg45 in the EXXER motif of TMH1 close to the binding site interacts with nitrate and further stabilizes the nitrate in the binding site. The tight binding of nitrate at the center of the transporter destabilizes the OF state and facilitates the conformational change to OC, where the extracellular gating distance reduces to ∼7Å (Fig. 4D). Nitrate dissociates from the central binding site and binds to Arg45, Lys164 and Tyr480. Our findings are consistent with the previous biochemical studies that the EXXER motif plays a crucial role in substrate translocation in PTR family, and that mutation of these hotspot residues results in loss of function.^12,13^ Parker and Newstead hypothesized that Lys164 and Glu476 form a salt bridge and mediate nitrate release to the cell.^13^ However, our simulations show that Glu476 forms polar contact with His356 to maintain the specificity of the binding site conformation for substrate binding, recognition and translocation. The nitrate engages with Lys164 and Tyr480, secondary gatekeeper residues that facilitate the opening of the intracellular gate and release of nitrate into the cells. Lys164 in the EXXER motif is conserved in POT family and mutation to alanine abrogates the transporter function.^12^ At this juncture, the extracellular surface is fully closed and the intracellular cavity partially opens as the gating residue distance increases from ∼4 Å to ∼8Å (Fig. 4E and F). (6) The nitrate ion diffuses down and forms polar contact with Arg183 (Fig. 4G) and enters into the interior of the cell as the intracellular cavity opens up to ∼16 Å to obtain the IF state (Fig. 4H).

**Fig. 4.**
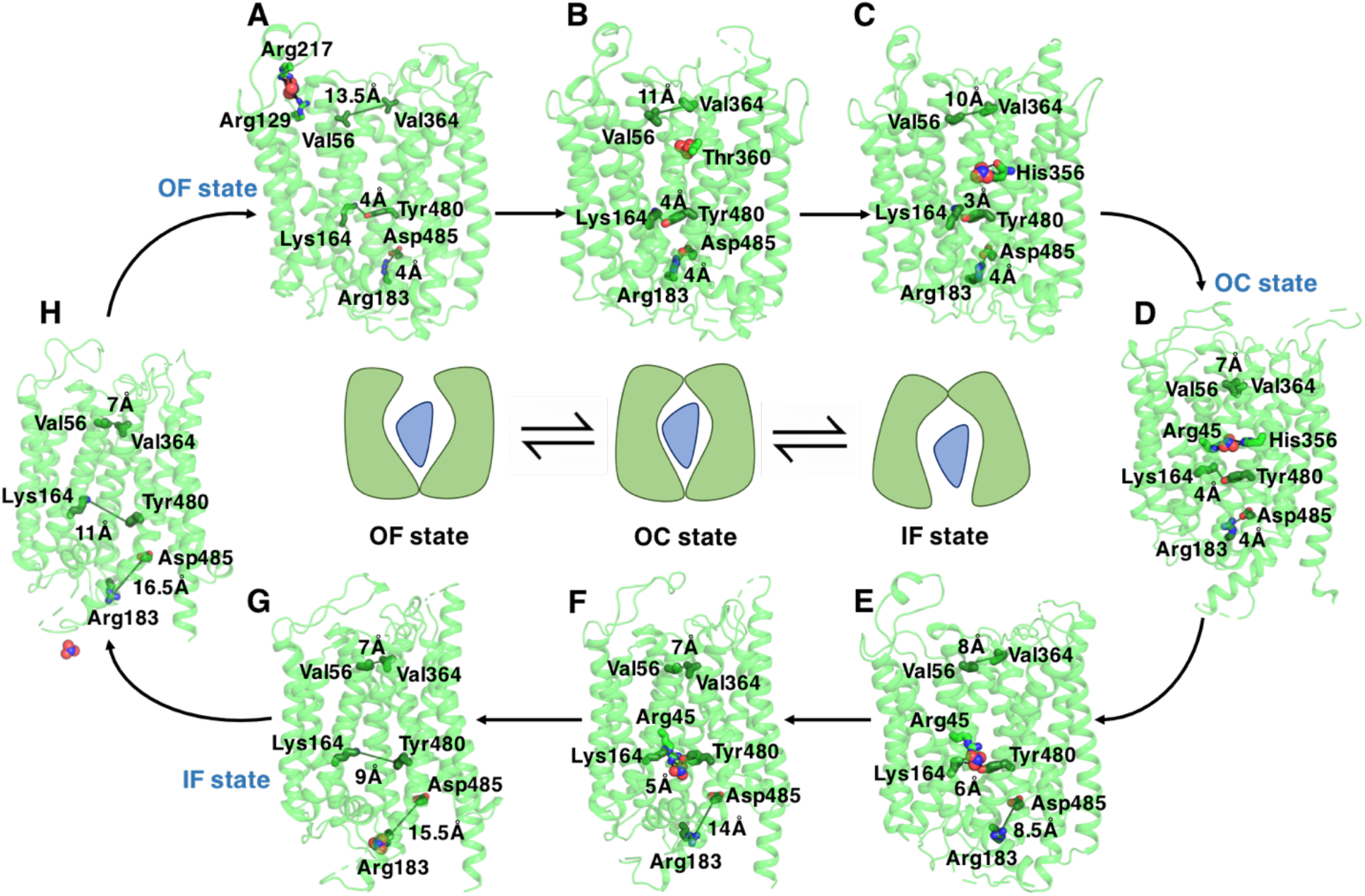
Nitrate transport mechanism in NRT1.1. Nitrate recognition, binding and transport driven conformational changes of NRT1.1 are shown here. Nitrate and interacting residues along the transport pore channel are shown in ball and stick representation (A-H). The gate opening was monitored by the distance between the residues Val56 (Cα)-Val364 (Cα), Lys164 (NZ)-Tyr480 (OH), and Arg183 (CZ)-Asp485 (CG), which were colored in green.

## CONCLUSIONS

Using molecular dynamics simulations in conjunction with Markov state models, we uncovered the molecular mechanisms of post-translation modification induced allostery in functional mechanism, conformational transition to intermediate states, and nitrate translocation in NRT1.1 at atomistic detail. Phosphorylation actively modifies the conformational landscape of the transporter and regulates the dynamics of intermediate states to function as highly efficient transporter. However, UnpNRT1.1 has multiple energy minima bridged by complex intermediate transition states. Further, phosphorylation enhances the dynamic coupling of distal residues and alters the transport properties of NRT1.1. Our kinetic data reveals that pNRT1.1 nitrate uptake rate is larger than UnpNRT1.1. The calculated free energy value for different states shows that nitrate interacts strongly with OC state in both pNRT1.1 and UnpNRT1.1, indicating that OC states act as an intermediate state to interconvert the conformation between OF and IF (Fig. S7). The umbrella sampling results demonstrate the functional relevance of the NRT1.1 dimer. The phosphorylation on Thr101 decouples motion between the two subunits, which behaves as an independent monomer. Taken together, our investigations to understand the role of phosphorylation reveal important regulatory mechanisms for nitrate uptake transporter. The recognition of nitrate ions also leads to signaling function in NRT1.1. However, further investigations are required to understand how nitrate translocation and sensing are coupled, which enables plants to survive in an environment with fluctuating nitrate availability.

In the past 50 years, the application of nitrogenous fertilizers has significantly increased crop productivity and alleviated world hunger. However, the production of nitrogenous fertilizers consumes ∼1% of global energy, and more than 50% of N resources is lost to the environment. Therefore, it is crucial to enhance the NUE for sustainable agriculture to increase the crop yield. Engineering crops to uptake more nitrogen resources are a growing concern in modern agriculture. Our study could potentially help plant community to understand the structure based functional mechanism of nitrate uptake transporter as a major step forward to reduce nitrogen pollution.

## Supporting information

Supplementary Images, Tables and Results

## ASSOCIATED CONTENT

### Supplementary Materials

All data needed to evaluate the conclusions are present in the paper and/or the Supplementary Materials. Additional data related to this paper may be requested from the authors. The Supplementary Materials contains the following:

Fig. S1. Top pathways of conformational transitions from IF to OF in UnpNRT1.1 and pNRT1.1 monomer.

Fig. S2. Kinetics of nitrate translocation in UnpNRT1.1 and pNRT1.1 monomer.

Fig. S3. Solvent-accessible surface area (SASA) values of Thr101 and its vicinity in IF, OC, and OF state.

Fig. S4. Structural rearrangements upon phosphorylation in the N-terminal bundle helices.

Fig. S5. Dynamic cross-correlation matrix (DCCM) analysis for all alpha carbon atoms in UnpNRT1.1 and pNRT1.1 monomer.

Fig. S6. Dynamic cross-correlation matrix (DCCM) analysis for all alpha carbon atoms in UnpNRT1.1 and pNRT1.1 dimer.

Fig. S7. Free energy of nitrate ion interacting with NRT1.1 under (A) OF, (B) OC, and (C) IF state.

Fig. S8. Implied timescale plots.

Fig. S9. GMRQ score with increasing number of iterations for UnpNRT1.1 and pNRT1.1 monomer.

Table S1. MSM hyperparameters selected by genetic algorithm.

## AUTHOR INFORMATION

### Author Contributions

D.S. conceived and supervised this study. B.S. and J.F. performed simulations and analyzed the simulation data. B.S. and J.F. wrote the manuscript with inputs from D.S.

### Funding Sources

D.S. acknowledges funding support from the New Innovator Award from the Foundation for Food and Agriculture Research, NSF Early Career Award (MCB 1845606) and Institute for Sustainability, Energy and Environment at University of Illinois. J.F. was supported by Chia-chen Chu Fellowship and Harry G. Drickamer Graduate Research Fellowship from University of Illinois, Urbana−Champaign, IL.

## ABBREVIATIONS

N: Nitrogen
NUE: nitrogen use efficiency
LATS: low-affinity transporter system
HATS: high-affinity transporter system
TM: transmembrane helix
CBL: Calcineurin-B Like
CIPK23: CBL-Interacting serine/threonine Protein Kinase
POT: proton-dependent oligopeptide transporter family
MFS: major facilitator superfamily
OF: outward-facing
OC: occluded
IF: inward-facing
UnpNRT1.1: unphosphorylated NRT1.1
pNRT1.1: phosphorylated NRT1.1
AtNRT1.1: *Arabidopsis thaliana* NRT1.1
MD: molecular dynamics
PDB: Protein Data Bank
POPC: phosphatidylcholine
PBC: periodic boundary conditions
PME: particle mesh Ewald
cMD: conventional MD
aMD: accelerated MD
MSM: Markov state model
tICA: time-lagged independent component analysis
MFPT: mean first passage time
PMF: potential of mean force
GMRQ: generalized matrix Raleigh quotient
dRMSD: distance root-mean-square displacement
TPT: transition path theory

## Notes

### Competing Interest Statement

The authors have declared no competing interest.

