## Supplementary Images, Tables and Results for "Atomistic Insights Into The Mechanism of Dual Affinity Switching In Plant Nitrate Transporter NRT1.1"

Fig. S1. Top pathways of conformational transitions from IF to OF in UnpNRT1.1 and pNRT1.1 monomer.
Fig. S2. Kinetics of nitrate translocation in UnpNRT1.1 and pNRT1.1 monomer.
Fig. S3. Solvent-accessible surface area (SASA) values of Thr101 and its vicinity in IF, OC, and OF state.
Fig. S4. Structural rearrangements upon phosphorylation in the N-terminal bundle helices.
Fig. S5. Dynamic cross-correlation matrix (DCCM) analysis for all alpha carbon atoms in UnpNRT1.1 and pNRT1.1 monomer.
Fig. S6. Dynamic cross-correlation matrix (DCCM) analysis for all alpha carbon atoms in UnpNRT1.1 and pNRT1.1 dimer.
Fig. S7. Free energy of nitrate ion interacting with NRT1.1 under (A) OF, (B) OC, and (C) IF state.
Fig. S8. Implied timescale plots.
Fig. S9. GMRQ score with increasing number of iterations for UnpNRT1.1 and pNRT1.1 monomer.
Table S1. MSM hyperparameters selected by genetic algorithm.

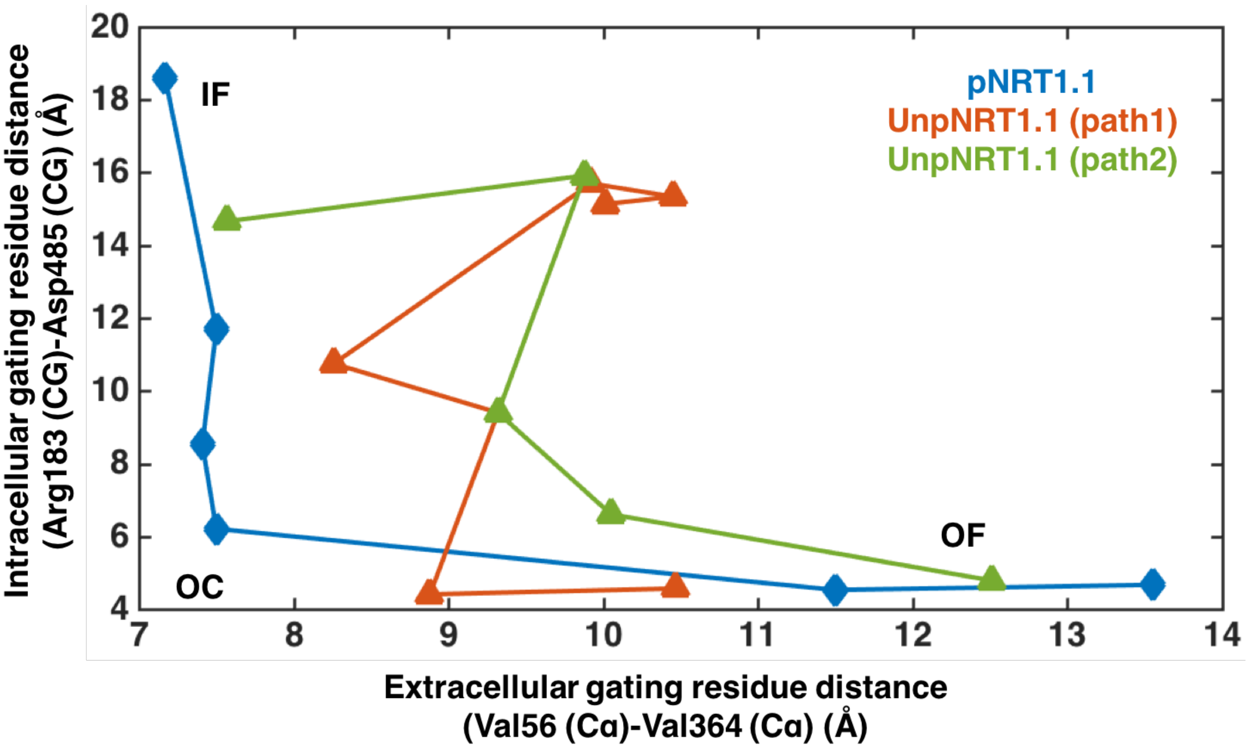


**Fig. S1. Top flux pathways of conformational transitions from IF to OF in UnpNRT1.1 and pNRT1.1 monomer.** The intracellular and extracellular gating opening was monitored by the distance between the residues Val56 (Cα)-Val364 (Cα) and Arg183 (CZ)-Asp485 (CG), respectively.

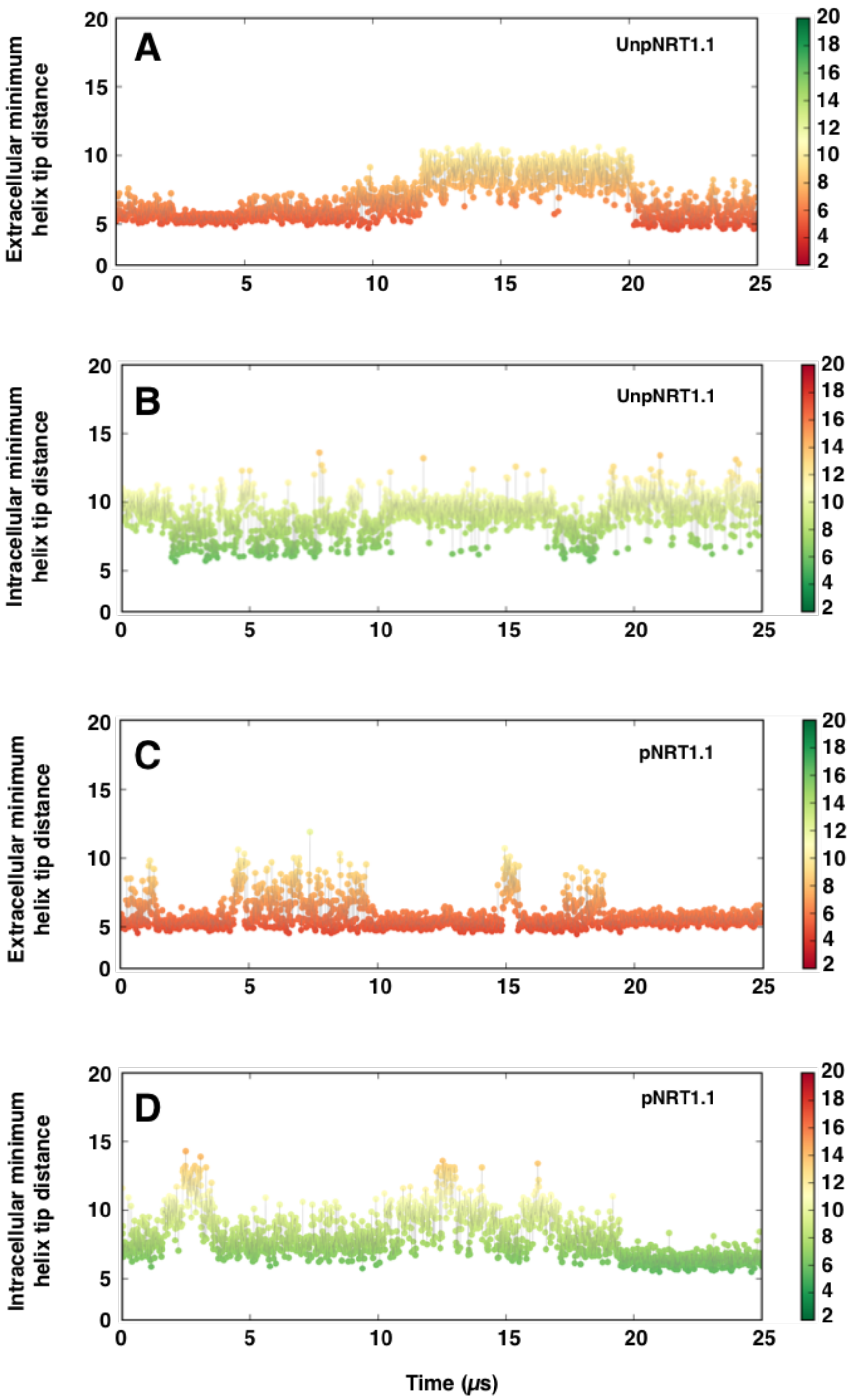


**Fig. S2. Kinetics of nitrate translocation in UnpNRT1.1 and pNRT1.1 monomer.** The kinetics plots were obtained using Kinetic Monte Carlo, which generates long time scale synthetic trajectories based on the MSM transition probability matrix. The color bar indicates the extent of opening and closing of the extracellular **(A, C)** and intracellular **(B, D)** gates. We generated extensive Kinetic Monte Carlo trajectories for a total of 25 μs for both UnpNRT1.1 and pNRT1.1 monomer.

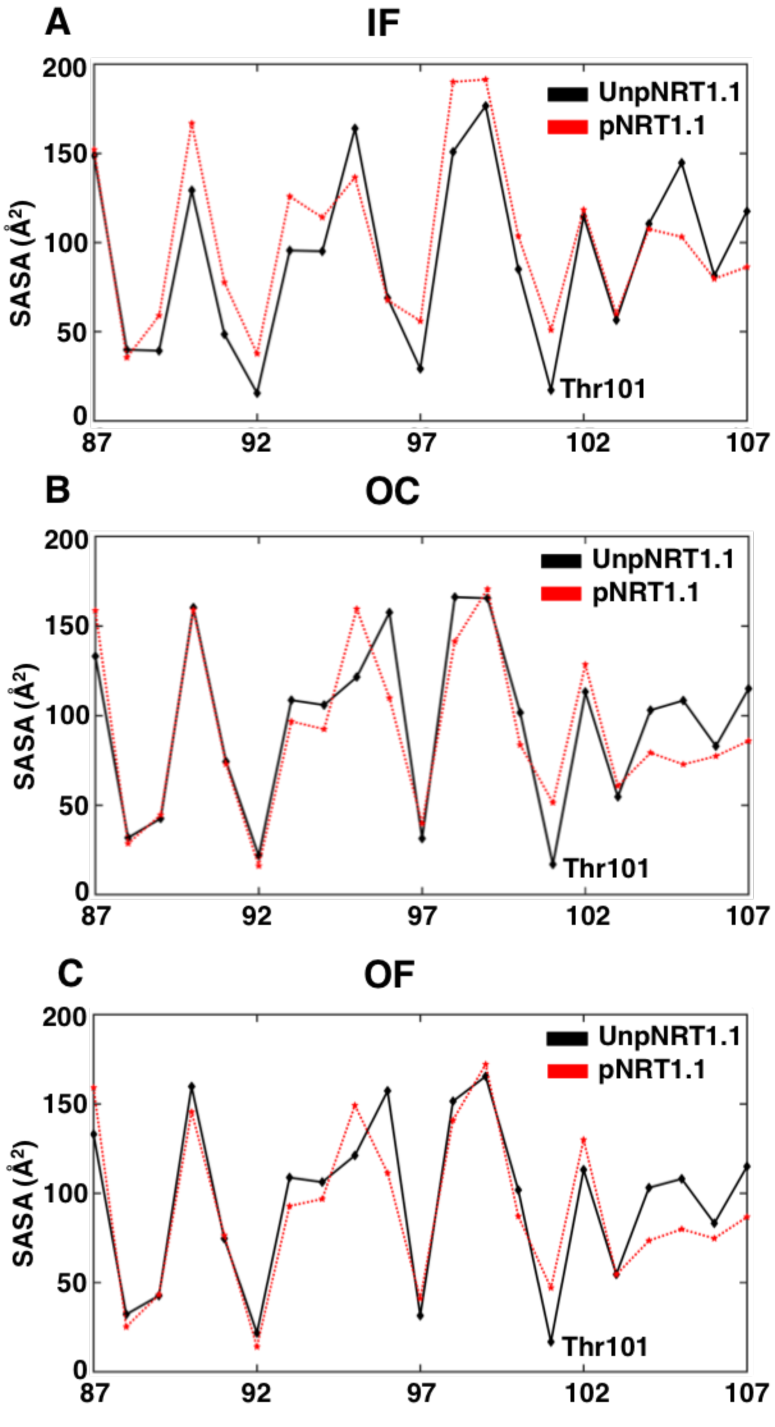


**Fig. S3. Solvent-accessible surface area (SASA) values of Thr101 and its vicinity in (A) IF, (B) OC, and (C) OF state.** Comparison between SASA values in the UnpNRT1.1 and pNRT1.1 monomers under three different conformational states suggests the exposure of Thr101 upon phosphorylation, with an increase of SASA values around ~ 30-40 Å$^2$.

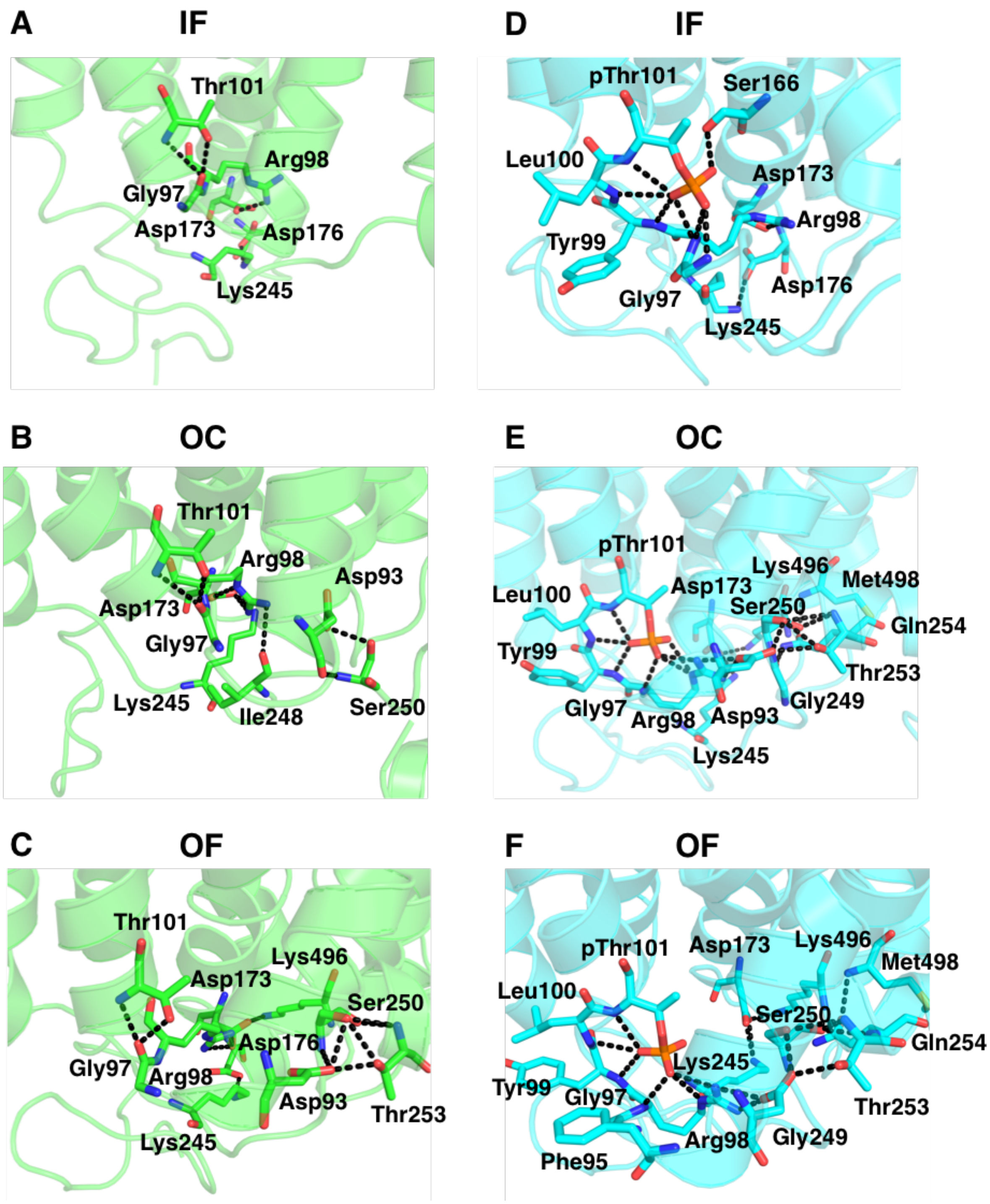


**Fig. S4. Structural rearrangements upon phosphorylation in the N-terminal bundle helices**. Interactions between Thr101 and neighboring residues are shown for UnpNRT1.1 **(A-C)** and pNRT1.1 **(D-F)**. Once phosphorylated, pThr101 forms an extensive network of polar interactions to facilitate the closure of intracellular gate.

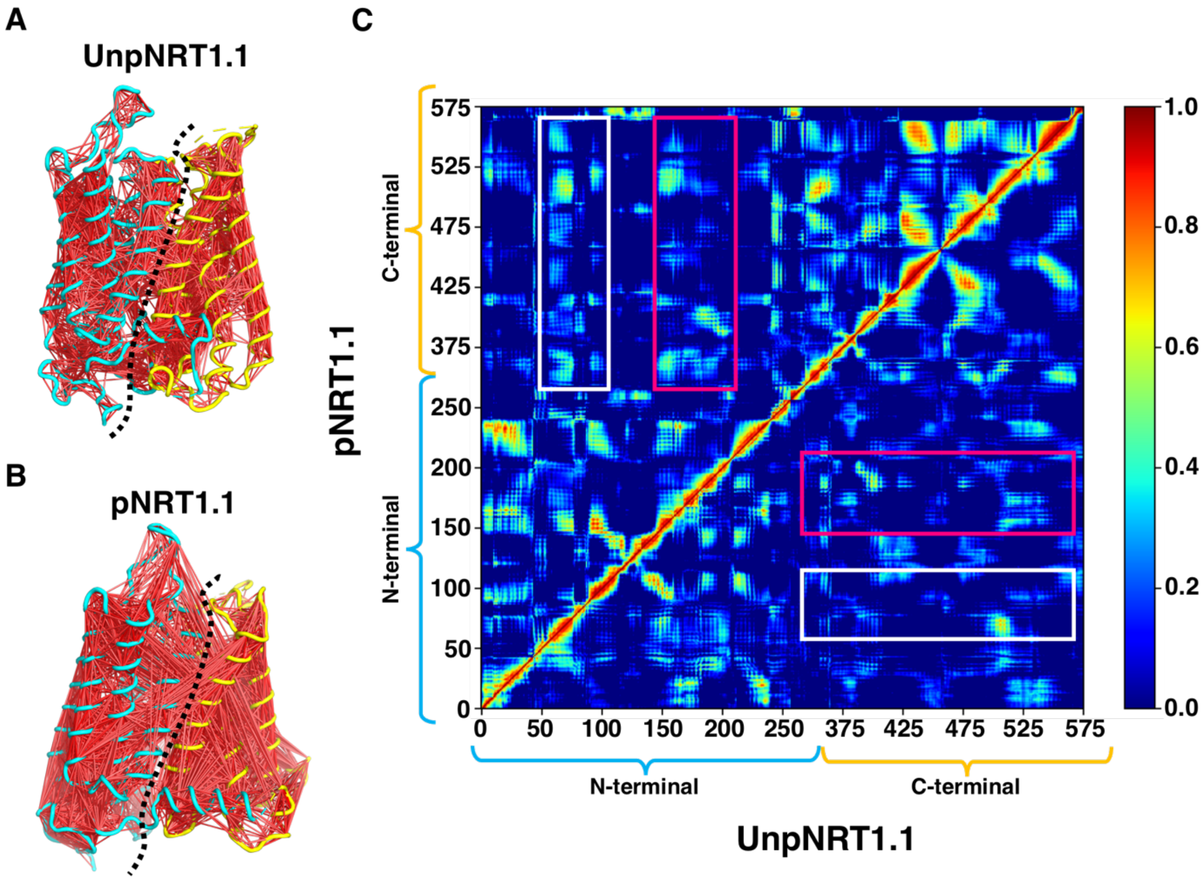


**Fig. S5. Dynamic cross-correlation matrix (DCCM) analysis for all alpha carbon atoms in UnpNRT1.1 (A, C) and pNRT1.1 monomer (B, C).** The N and C-terminal domains are colored in cyan and yellow, respectively (A, B). Red lines indicate strong dynamic coupling between two residues (cross-correlation coefficient > 0.6) (A, B). The matrix plot shows the correlation between any two residues, ranging from 0 to 1 as indicated by the color bar (C). The closer the value to 1, the stronger the dynamic coupling between two residues. The boxed regions show the regions that strongly coupled in pNRT1.1 but weakly coupled in UnpNRT1.1 (C). These results suggest an enhanced dynamic coupling between N and C domains upon phosphorylation.

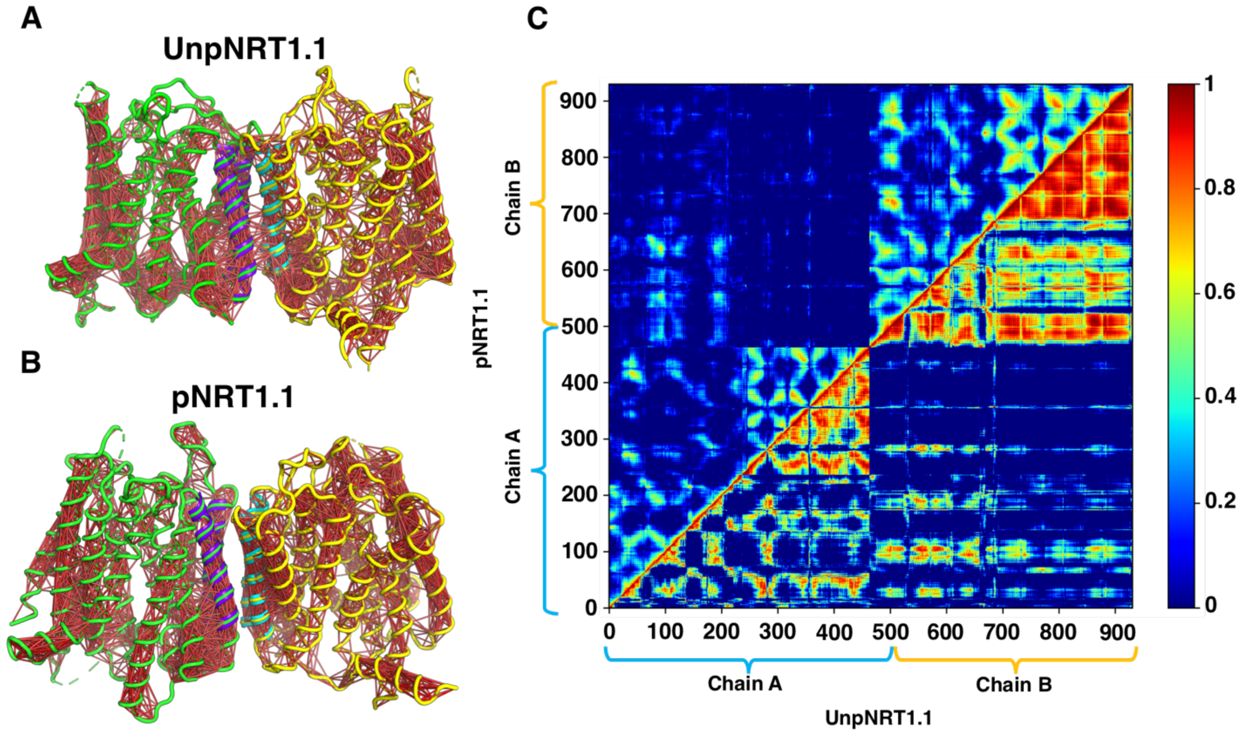


**Fig. S6. Dynamic cross-correlation matrix (DCCM) analysis for all alpha carbon atoms in UnpNRT1.1 (A, C) and pNRT1.1 dimer (B, C).** Chain A and Chain B are colored in green and yellow, respectively (A, B). The dimer interface is colored in purple for Chain A and cyan for Chain B (A, B). Red lines indicate strong dynamic coupling between two residues (cross-correlation coefficient > 0.6) (A, B). The matrix plot shows the correlation between any two residues, ranging from 0 to 1 as indicated by the color bar (C). The closer the value to 1, the stronger the dynamic coupling between two residues. We observe stronger dynamic coupling between two monomers in UnpNRT1.1 compared with pNRT1.1 dimer (C).

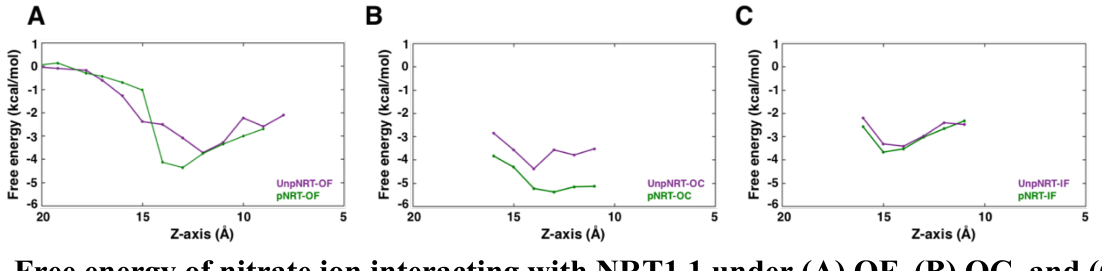


**Fig. S7. Free energy of nitrate ion interacting with NRT1.1 under (A) OF, (B) OC, and (C) IF state.** Comparison of free energy shows that nitrate exhibits stronger interactions with OC state in both UnpNRT1.1 and pNRT1.1.

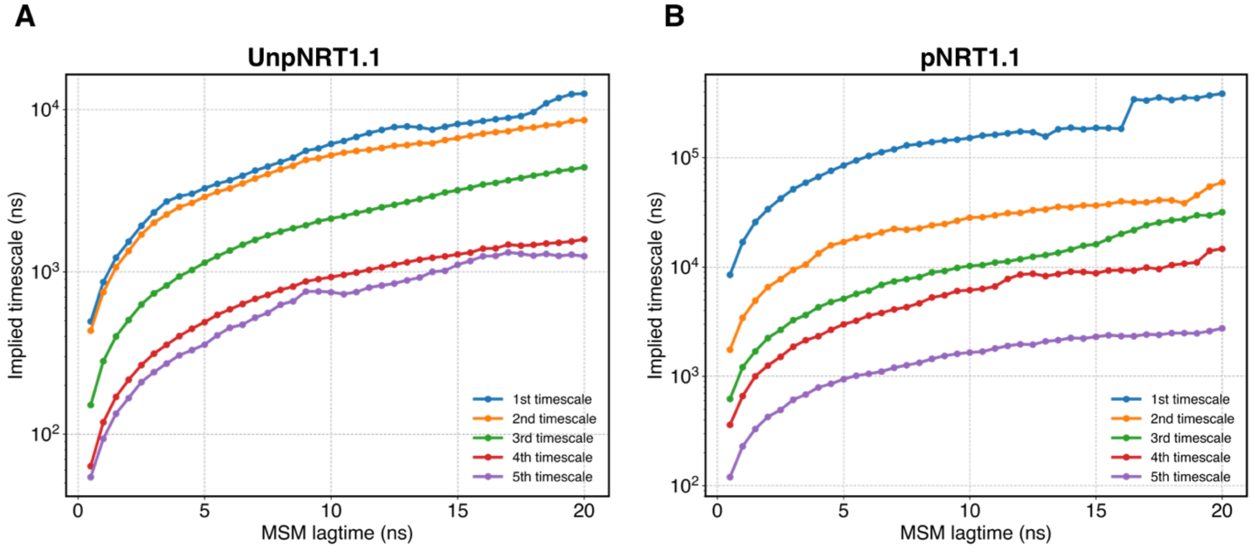


**Fig. S8. Implied timescale plots.** Five slowest timescales varying with different MSM lagtimes are shown for **(A)** UnpNRT1.1 and **(B)** pNRT1.1 monomer. The MSM lagtime was chosen as 14.3 ns such that the five slowest timescales converge.

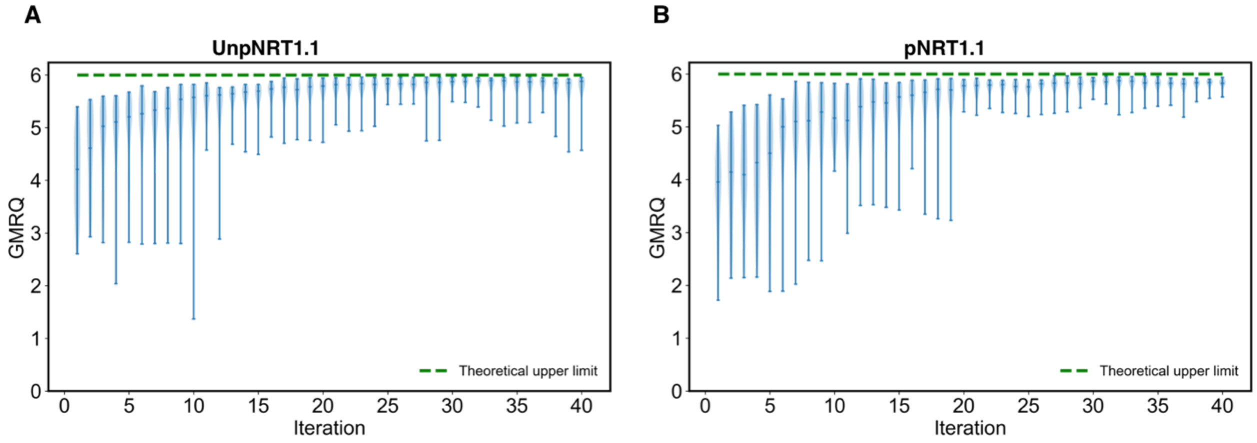


**Fig. S9. GMRQ scores with increasing number of iterations for (A) UnpNRT1.1 and (B) pNRT1.1 monomer.** Green, dashed lines indicate the theoretical upper limit for GMRQ score under given number of MSM timescales. Each violin plot shows the convergence of GMRQ scores over 40 iterations.

**Table S1. MSM hyperparameters selected by genetic algorithm.**

| Protein | Number of chosen distances | tICA components | tICA lag time (ns) | Number of clusters | Number of MSM timescales | MSM lag time (ns) | Best GRMQ |
|---|---|---|---|---|---|---|---|
| UnpNRT1.1 | 44 | 5 | 0.1 | 200 | 5 | 14.3 | 5.96 |
| pNRT1.1 | 42 | 5 | 0.1 | 200 | 5 | 14.3 | 5.97 |
